# LCN2+ Astrocytes Mark a Pro-inflammatory Astrocyte State Associated with Secondary Brain Injury After Cerebral Venous Thrombosis

**DOI:** 10.64898/2026.11.04.764144

**Authors:** Ling Kui, Si-Wei Zhou, Yin-Ming Jiao, Rong Zhang, Ling-Bin Liu, Ke-Xu Xiong, Guo-Yun Wang, Yan-Qing Zhou

**Affiliations:** Shenzhen Bao’an Chinese Medicine Hospital, Shenzhen, Guangdong, China; Shenzhen Qianhai Shekou Free Trade Zone Hospital, Shenzhen, Guangdong, China; Shenzhen Key Laboratory of Viral Oncology, Clinical Innovation and Research Centre (CIRC), Shenzhen Hospital of Southern Medical University, Shenzhen, Guangdong, China; Shenzhen School of Clinical Medicine, Southern Medical University, Shenzhen, Guangdong, China; Research and Development Department, Shenzhen Mlay Intelligent Technology Co., Ltd., Shenzhen, Guangdong, China; Guangdong Medical University, Dongguan 523808, Guangdong, China

## Abstract

Cerebral venous thrombosis (CVT) is a cerebrovascular disorder with unique pathophysiological characteristics. Although the mortality rate during the acute phase has declined in recent years, the cellular mechanisms underlying secondary brain injury and long-term neurological dysfunction after CVT remain poorly understood [1,2]. In this study, we performed single-cell RNA sequencing of brain tissues from a murine CVT model to systematically characterize the cellular landscape following CVT and identified a reactive astrocyte state characterized by high expression of Lcn2. This astrocyte subset was markedly enriched after CVT. Notably, despite the absence of significant changes in the overall proportion of astrocytes, this population exhibited enhanced inflammatory/cytokine response signatures, ferroptosis-related stress responses, and vascular-immune microenvironment remodeling programs, suggesting that the major alteration of astrocytes after CVT is driven primarily by state transition rather than cellular expansion.

Pseudotime trajectory analysis further demonstrated that Lcn2⁺ astrocytes were predominantly localized at the late stage of the trajectory and displayed sustained upregulation of multiple inflammation- and stress-associated genes, supporting their characterization as a late-stage reactive astrocyte state under CVT conditions. Cell-cell communication analysis revealed extensive predicted interactions between this subset and myeloid cells, vascular-associated cells, as well as neuronal/synaptic populations, indicating its potential involvement in dynamic remodeling of the lesion microenvironment.

Cross-model integration analysis further revealed the presence of Lcn2-high astrocytes in the middle cerebral artery occlusion (MCAO) model; however, their expression breadth and intensity were generally weaker than those observed in CVT, suggesting that this cellular state may represent a common reactive framework following cerebrovascular obstruction, while being particularly pronounced in CVT. Furthermore, genetic deletion of LCN2 alleviated CVT-induced brain tissue injury in vivo, providing functional evidence supporting the detrimental role of the LCN2-associated pathological state.

Collectively, our study identifies a distinct Lcn2-high reactive astrocyte state after CVT that is closely associated with inflammatory amplification, ferroptosis-related stress, and microenvironmental remodeling. These findings provide new cellular insights into the mechanisms underlying secondary brain injury after CVT and highlight potential targets for therapeutic intervention.

## Introduction

Cerebral venous thrombosis (CVT) is a relatively uncommon cerebrovascular disorder, yet its clinical consequences should not be underestimated [1,3]. Unlike arterial ischemic stroke, CVT is primarily driven by impaired venous drainage and sustained elevation of venous pressure, which may subsequently lead to cerebral venous congestion, disruption of the blood-brain barrier, cerebral edema, and hemorrhagic infarction [4–6]. Although advances in early diagnosis and anticoagulant therapy have substantially improved acute outcomes in patients with CVT, a considerable proportion of survivors continue to experience long-term neurological sequelae, including cognitive impairment, epilepsy, chronic headache, and neuropsychiatric symptoms [7–10].

Current understanding of secondary brain injury following CVT is still largely derived from conceptual frameworks established in arterial ischemic stroke. However, CVT differs fundamentally from arterial ischemia in terms of its hemodynamic basis, pattern of tissue injury, and inflammatory milieu. Therefore, its underlying cellular and molecular mechanisms cannot simply be regarded as a venous counterpart or mirror image of arterial ischemic injury [1,4]. In particular, although previous studies have shown that multiple brain-resident and infiltrating cell populations become activated and interact following CVT, a comprehensive understanding of microenvironmental remodeling at single-cell resolution remains lacking and requires further systematic investigation [11–13].

Astrocytes are essential for maintaining central nervous system homeostasis and contribute to metabolic support, synaptic regulation, preservation of blood-brain barrier integrity, and neuroimmune balance [14–16]. In recent years, advances in single-cell and spatial transcriptomics have substantially reshaped our understanding of astrocyte biology. Rather than constituting a homogeneous cell population, astrocytes can adopt diverse reactive states with distinct molecular features and functional orientations in stroke, traumatic injury, infection, and neurodegenerative diseases [17–19]. These states may exert context-dependent effects: some restrict the spread of tissue damage, whereas others amplify inflammation, compromise vascular barrier integrity, and aggravate neuronal injury [18]. By contrast, whether astrocytes undergo disease-specific state remodeling in the context of CVT, and how such states contribute to secondary brain injury, remain largely unknown.

Lipocalin-2 (LCN2) is a stress-inducible secreted protein that has attracted increasing attention as a molecular link among inflammatory amplification, disruption of iron homeostasis, and oxidative injury [20]. In models of arterial ischemic stroke, intracerebral hemorrhage, and other forms of acute brain injury, LCN2 has been shown to promote inflammatory responses, impair blood-brain barrier integrity, and increase cellular susceptibility to iron-dependent oxidative stress and lipid peroxidation [21–23]. However, the role of LCN2 in CVT has not been systematically defined. In particular, within the distinctive pathological context of CVT, which is characterized by venous congestion, increased vascular barrier vulnerability, and sustained inflammatory activation, several important questions remain unanswered: whether LCN2 expression is driven by a specific cellular state, whether it acts as a key amplifier of CVT-associated brain injury, and which molecular programs are shared with, or distinct from, those observed in other cerebrovascular injury models.

Notably, CVT and arterial cerebral ischemia do not represent two entirely unrelated forms of brain injury [2,11]. Both can induce tissue hypoxia-ischemia, sterile inflammatory activation, and glial responses, and may therefore share certain injury-response programs. Nevertheless, the distinctive features of CVT, including elevated venous pressure, extensive venous congestion, increased blood-brain barrier vulnerability, and hemorrhagic components, may give rise to glial states that differ from those observed in arterial ischemia models such as middle cerebral artery occlusion (MCAO) [2,13,24]. In other words, CVT-associated cellular programs are likely to exhibit both commonality and specificity. On the one hand, they may share a general post-injury reactive framework with MCAO; on the other hand, they may display shifts in intensity, spatial distribution, or molecular composition that more closely reflect the characteristics of venous brain injury. Directly comparing this “shared but distinct” relationship may not only enable a more precise definition of the cellular pathology of CVT, but also prevent reactive astrocytes arising from different forms of cerebrovascular injury from being indiscriminately classified as a single state.

Accordingly, in the present study, we performed single-cell RNA sequencing of brain tissues from a murine CVT model to systematically characterize the cellular landscape following CVT, with a particular focus on astrocyte-state heterogeneity. We identified a reactive astrocyte state characterized by high expression of Lcn2 and systematically characterized its molecular programs, dynamic evolution, and microenvironmental interactions through pathway enrichment, pseudotime trajectory, and cell-cell communication analyses. We further incorporated an MCAO model for cross-model integration and comparison to determine the extent to which this state represents a shared response to cerebrovascular injury and the extent to which it is particularly prominent in CVT. In parallel, in vivo experiments using LCN2-deficient mice were conducted to functionally assess the contribution of LCN2 to secondary brain injury following CVT. Through these analyses, we sought to address several key questions: whether CVT induces a pathological astrocyte state characterized by high Lcn2 expression; where this state is positioned along the trajectory of astrocyte-state evolution; whether it is closely associated with inflammatory amplification, ferroptosis-related stress, and vascular-immune microenvironment remodeling; and whether it represents a general response to cerebrovascular injury or a pathologically distinct state that is more pronounced in CVT.

Overall, this study aims to redefine the process of brain microenvironment remodeling following CVT at single-cell resolution and proposes a more mechanistically oriented framework: rather than merely inducing nonspecific glial activation, CVT promotes the emergence of an Lcn2-high reactive astrocyte state associated with inflammation, iron-related stress, and reorganization of cell-cell communication networks. Cross-model comparison of the shared and disease-specific features of this state, together with in vivo functional validation, may not only deepen our understanding of the mechanisms underlying secondary brain injury after CVT, but also provide a theoretical basis for precision therapeutic strategies targeting LCN2-associated astrocyte states.

## Results

### Single-cell transcriptomic profiling reveals the cellular landscape and microenvironmental remodeling after CVT

To systematically characterize alterations in the cellular composition and microenvironment of the brain following cerebral venous thrombosis (CVT), we performed single-cell RNA sequencing of brain tissues obtained from mice subjected to CVT modeling (n = 6) and sham-operated controls (n = 6). Following stringent quality control, data integration, and dimensionality reduction, UMAP visualization resolved multiple major cell populations, including astrocytes, microglia, infiltrating macrophages, neurons, oligodendrocytes and oligodendrocyte precursor cells, endothelial cells, pericytes, fibroblasts, choroid plexus cells, erythrocytes, and infiltrating lymphocytes (Fig. 1A). Quality control metrics and canonical marker gene expression supporting cell-type annotation are shown in Supplementary Fig. 1.These findings indicate that the single-cell transcriptomic dataset reliably captured the cellular composition of brain tissue following CVT.

**Figure 1.**
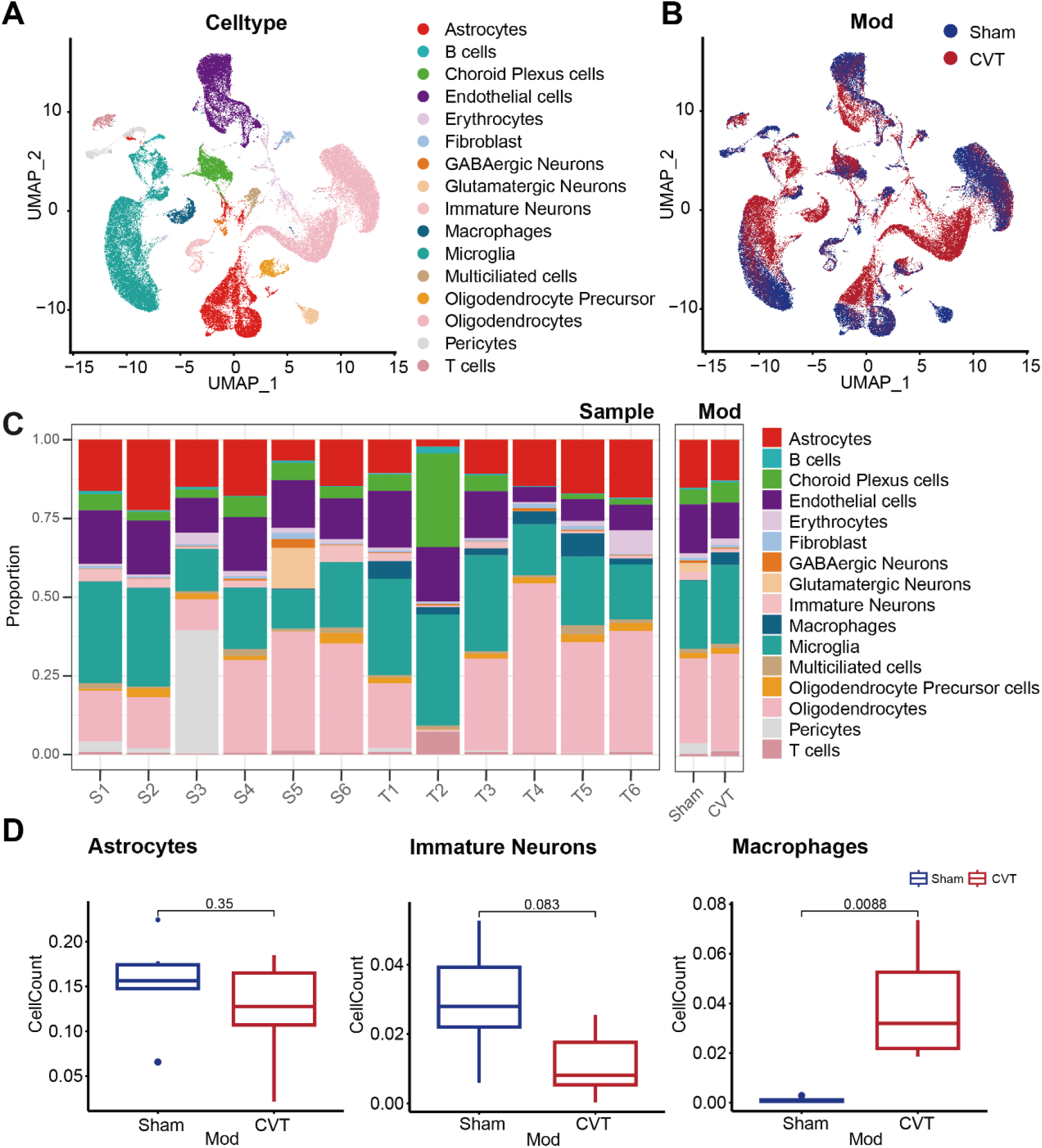
Single-cell landscape and cell-type composition of the CVT-injured brain. (A) UMAP visualization of integrated single-cell RNA-seq data showing major cell types identified in mouse brain tissues, including astrocytes, microglia, macrophages, neurons, oligodendrocyte lineage cells, endothelial cells, pericytes, and other neural and non-neural populations. (B) UMAP projection colored by experimental condition, illustrating the distribution of cells from sham (S) and CVT (T) groups across the transcriptional landscape. (C) Stacked bar plots showing the relative proportions of annotated cell types in individual samples from sham (S1-S6) and CVT (T1-T6) groups, as well as the aggregated proportions for each condition. (D) Box plots comparing the relative abundance of representative cell populations between sham and CVT groups, including immature neurons, macrophages, and astrocytes. P values are indicated above each comparison.

Building on this cellular annotation, we compared the distribution of cells between the sham control (S) and CVT (T) groups. Although cells from the two groups partially overlapped within the global transcriptional space, UMAP visualization revealed marked shifts in multiple regions in the CVT group (Fig. 1B), suggesting that CVT induces substantial transcriptomic remodeling rather than merely reflecting cell loss or technical variation. Further sample-level quantification of the relative abundance of each cell type demonstrated pronounced alterations in the cellular composition of the brain following CVT, characterized by an increased proportion of myeloid cells, together with varying changes in the proportions of neurons and several glial cell populations (Fig. 1C).

To determine the statistical significance of these alterations, we performed between-group comparisons of representative cell populations. Compared with the sham control group, the CVT group exhibited a significant increase in macrophages and a marked reduction in immature neurons (Fig. 1D), indicating enhanced inflammatory cell recruitment and impaired neuronal homeostasis following CVT. Notably, despite the pronounced inflammatory response within the brain, the overall proportion of astrocytes did not differ significantly between the two groups (Fig. 1D). Collectively, these findings demonstrate that CVT does not merely alter cellular composition but instead drives systematic remodeling of the brain microenvironment toward an inflammation-dominated cellular ecosystem.

### CVT induces the emergence of an Lcn2⁺ reactive astrocyte state characterized by inflammatory and cytokine-response programs

To further characterize astrocyte heterogeneity following CVT, we isolated astrocytes from the integrated single-cell dataset and performed subclustering analysis. UMAP visualization resolved astrocytes into multiple transcriptionally distinct subclusters, indicating substantial functional heterogeneity within the astrocyte lineage (Fig. 2A). When stratified by experimental group, several astrocyte subclusters were markedly enriched in the CVT group but were relatively underrepresented in the sham group, suggesting that CVT drives pronounced transitions in astrocyte states (Fig. 2A).

**Figure 2.**
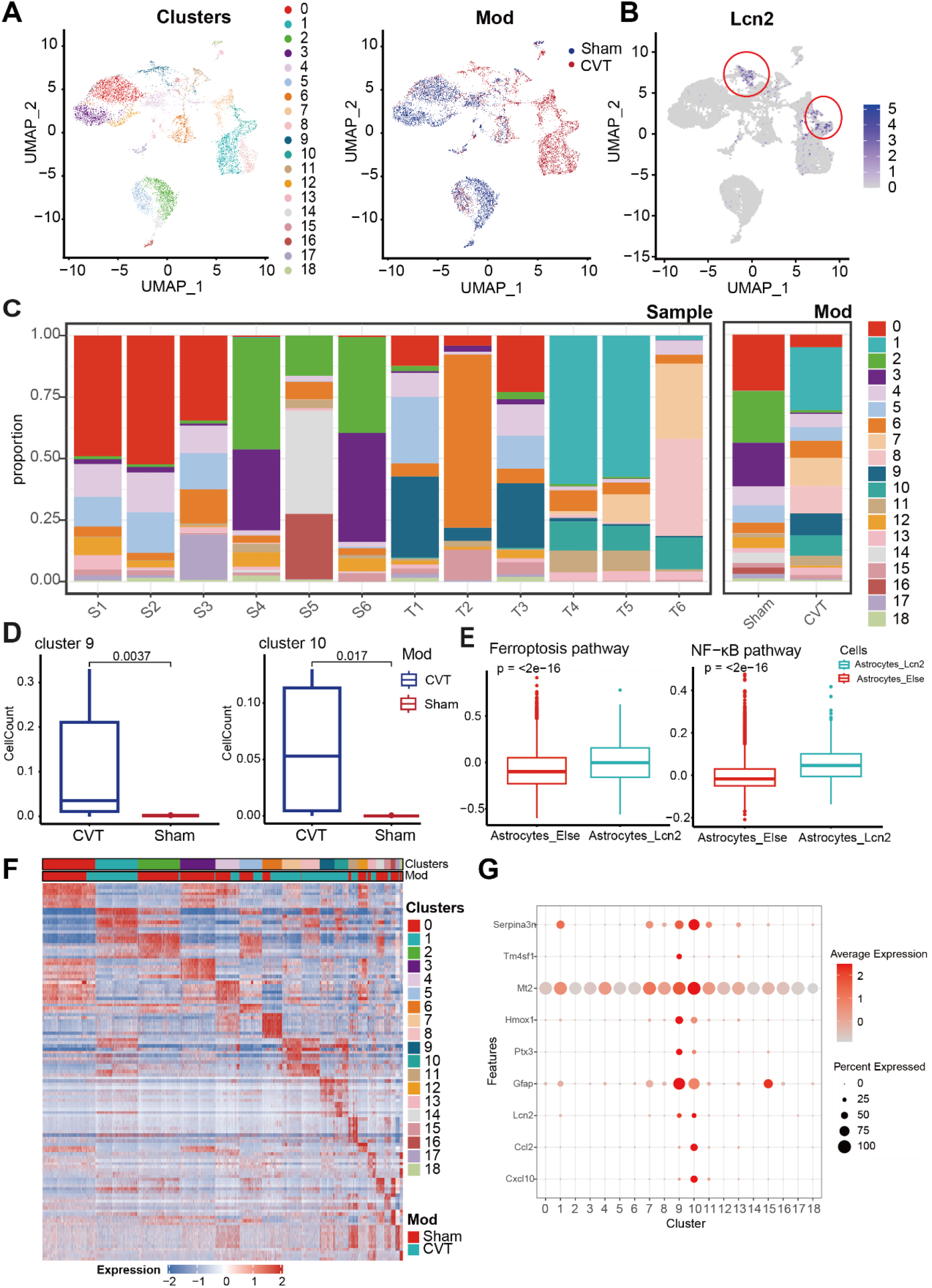
A distinct LCN2+ astrocyte state emerges following cerebral venous thrombosis. (A) UMAP visualization of astrocyte subclusters identified by unsupervised clustering, colored by cluster identity (left) and experimental condition (right; sham vs. CVT). (B) Feature plot showing Lcn2 expression across astrocyte clusters, highlighting discrete LCN2+ astrocyte subpopulations (red circles). (C) Stacked bar plots depicting the relative proportions of astrocyte clusters across individual samples and aggregated by experimental condition. (D) Box plots comparing the proportions of representative LCN2-enriched astrocyte clusters (clusters 9 and 10) between sham and CVT groups. P values are indicated. (E) Box plots showing gene set enrichment scores for ferroptosis and NF-κB signaling pathways in LCN2+ versus LCN2-astrocytes. (F) Heatmap of scaled expression values for selected differentially expressed genes across astrocyte clusters, annotated by cluster identity and experimental condition. (G) Bubble plot showing the expression of characteristic genes in Lcn2+ astrocytes.

FeaturePlot analysis of key gene expression further revealed a pronounced subcluster-specific distribution of Lcn2 within astrocytes. Lcn2 expression was restricted to a small number of cell populations, forming a spatially discrete Lcn2⁺ astrocyte subset on the UMAP (Fig. 2B). The restricted expression pattern of Lcn2 across astrocyte subclusters was further confirmed by violin plot analysis, demonstrating preferential enrichment of Lcn2 expression within specific astrocyte populations (Fig. S2A). These Lcn2⁺ astrocytes were derived predominantly from CVT samples, whereas Lcn2 expression was minimal in the sham group, suggesting that Lcn2 upregulation is not a constitutive feature of astrocytes but rather represents a stress-responsive state induced by CVT.

To quantitatively assess changes in subcluster distribution, we performed sample-level statistical analysis of astrocyte subcluster composition. The Lcn2-enriched subclusters were significantly expanded in CVT mice, with clusters 9 and 10 accounting for markedly higher proportions in the CVT group than in the sham group (Fig. 2C,D). Consistent with these findings, sample-level quantification further demonstrated an increased abundance of Lcn2-high astrocytes following CVT (Fig. S2B). These findings supported the designation of clusters 9 and 10 as CVT-associated Lcn2⁺ astrocyte states.

Building on these findings, we further characterized the molecular features of this astrocyte state from a functional perspective. Pathway enrichment analysis revealed that Lcn2⁺ astrocytes were significantly enriched for ferroptosis-related pathways and exhibited activation of NF-κB signaling (Fig. 2E), suggesting an activated state characterized by inflammatory amplification and iron-dependent stress. Hallmark pathway scoring further showed that clusters 9 and 10, corresponding to the Lcn2⁺ astrocyte state, displayed among the highest activity levels across multiple inflammation- and cytokine-related pathways, including IL-2-STAT5 and IL-6-JAK-STAT3 signaling (Fig. 3A). As key regulatory axes of inflammatory amplification and stress-responsive transcription, these pathways suggest that Lcn2⁺ astrocytes exhibit heightened responsiveness to cytokine stimulation following CVT.

**Figure 3.**
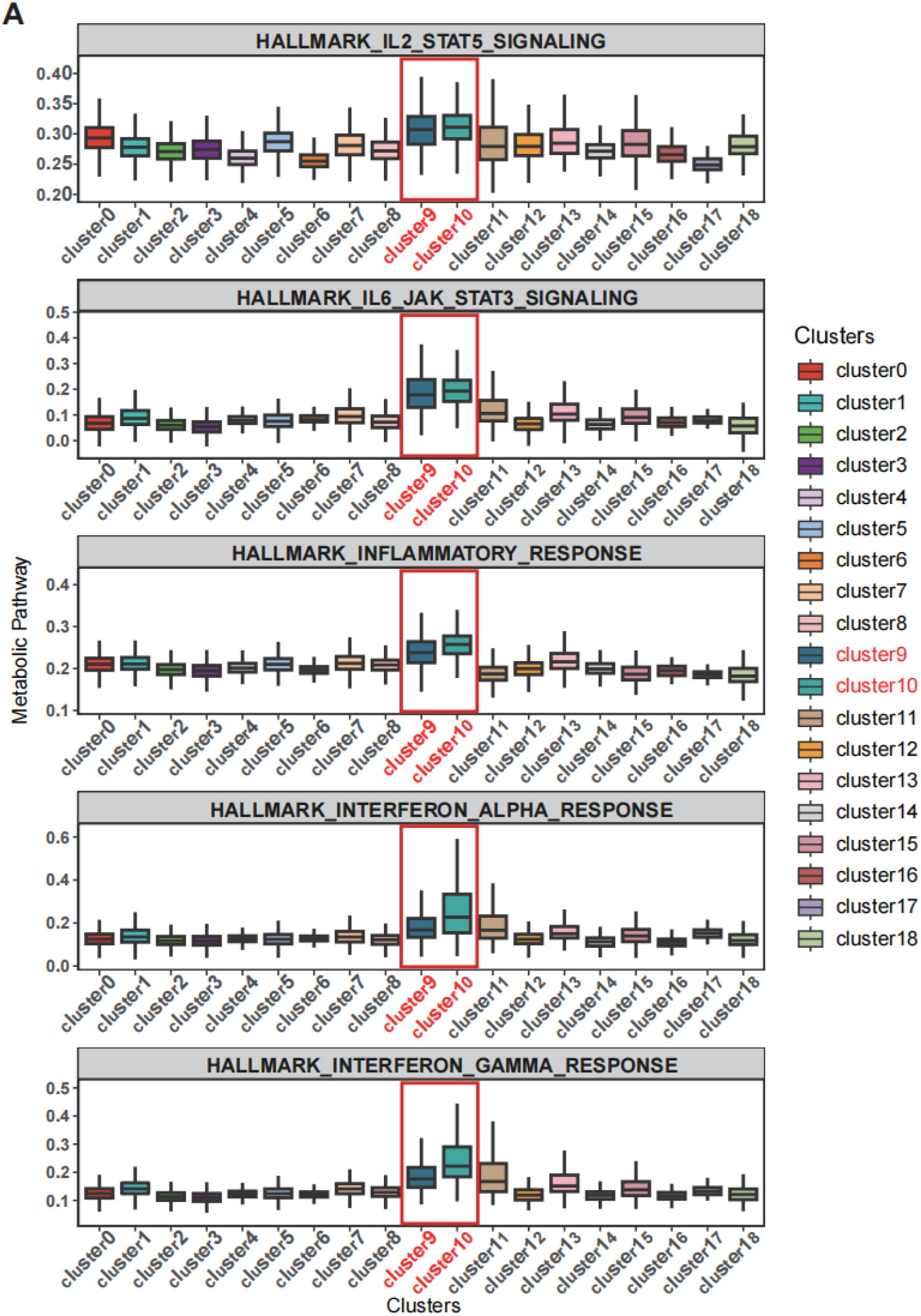
LCN2+ astrocytes display enhanced cytokine and inflammatory signaling activity. (A) Box plots showing Hallmark pathway enrichment scores across astrocyte clusters for IL-2-STAT5 signaling, IL-6-JAK-STAT3 signaling, inflammatory response, interferon-α response, and interferon-γ response. Clusters 9 and 10 (red boxes), corresponding to LCN2-enriched astrocyte populations, exhibit consistently elevated pathway activity compared with other astrocyte clusters.

In addition, the Lcn2-enriched subclusters showed markedly increased inflammatory and interferon-related signaling, including interferon-α and interferon-γ response programs (Fig. 3A). Notably, activation of these pathways was largely restricted to clusters 9 and 10 (Supplementary Table 1) and was not broadly observed across the other astrocyte subclusters, further highlighting the pronounced functional heterogeneity of astrocytes following CVT.

Consistent with these findings, correlation analysis placed Lcn2 at the center of a transcriptional module characterized by inflammatory responses, oxidative stress, and extracellular matrix remodeling, with its expression showing coordinated variation with multiple immune activation-associated genes (Fig. 2F). Hierarchical clustering further demonstrated that Lcn2⁺ astrocytes formed a transcriptionally coherent cell population that was largely restricted to CVT samples (Fig. 2G), further supporting its identity as a distinct disease-associated astrocyte state.

### CVT drives astrocyte transition toward an Lcn2-high reactive state along the pseudotime trajectory

To characterize the dynamic evolution of astrocyte states under CVT conditions, we performed Monocle-based pseudotime trajectory analysis of the astrocyte population. Astrocytes were distributed along a continuous state trajectory with multiple branching structures (Fig. 4A). Visualization according to experimental group showed that cells from the sham control group (S) were predominantly located at the early stage of the trajectory and along several homeostatic branches, whereas cells from the CVT group (T) were relatively enriched in the rightward and lower branches. These findings suggest that CVT drives astrocytes toward distinct reactive states.

**Figure 4.**
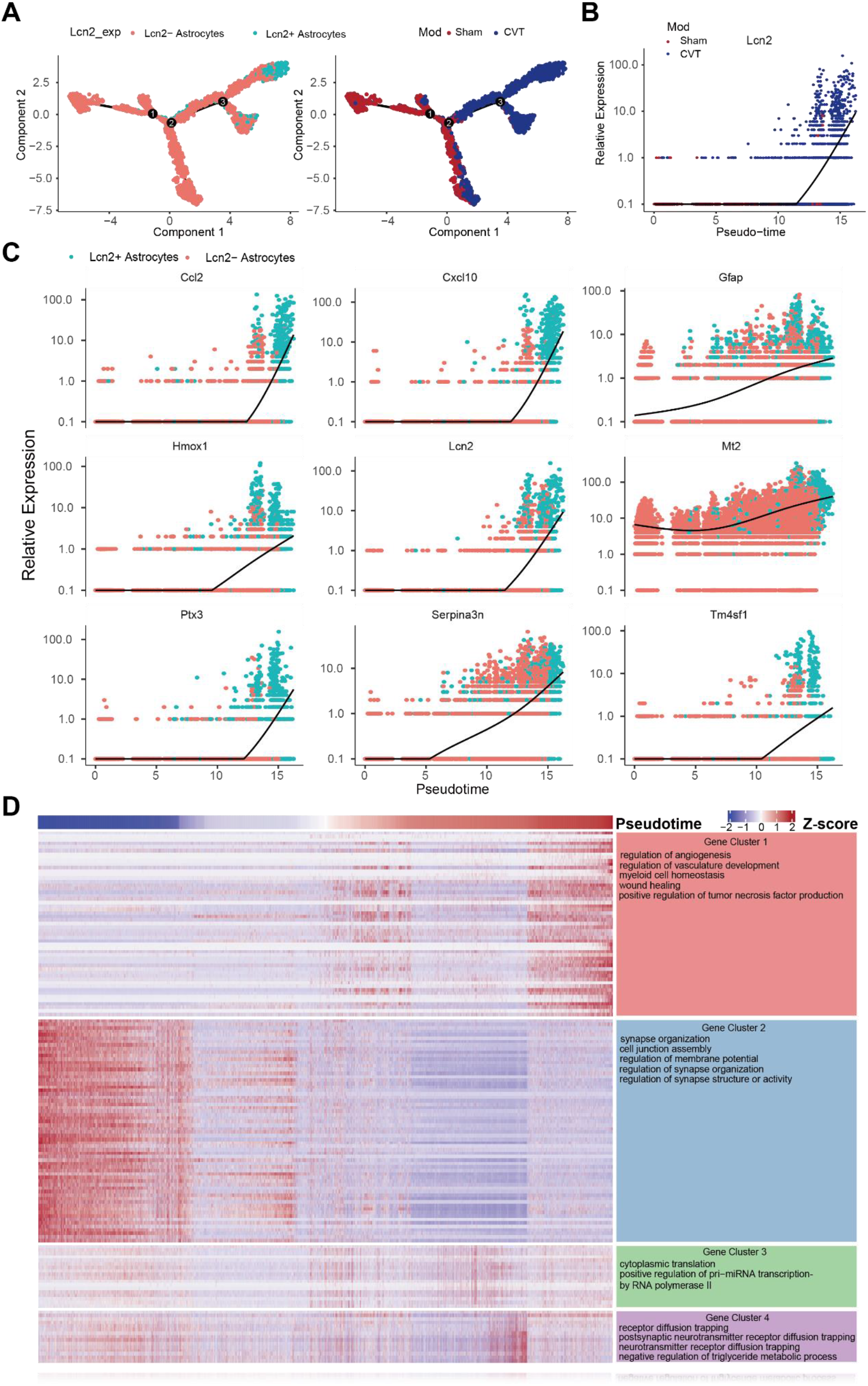
Pseudotime analysis reveals a progressive transition of astrocytes toward an Lcn2-high reactive state in CVT. (A) Monocle pseudotime trajectory plots, colored by *mod* (left) and by *Lcn2_exp* (right). (B) *Lcn2* expression gradually increases along pseudotime, with the upregulation predominantly observed in T group cells. (C) Expression dynamics of representative genes associated with Lcn2+ astrocytes along pseudotime. (D) Heatmap of dynamic genes showing that late pseudotime stages are characterized by activation of inflammatory, stress-response, and reactive gliosis-related transcriptional programs.

Further stratification according to Lcn2 expression revealed that Lcn2⁺ astrocytes were predominantly localized to the late stages and terminal branches of the trajectory, whereas Lcn2⁻ cells were broadly distributed across the early and intermediate stages (Fig. 4A). This distribution suggests that high Lcn2 expression marks a late-stage state during astrocyte-state progression. Consistent with this observation, Lcn2 expression remained low during the early pseudotime stages but increased markedly toward the late stages (Fig. 4B). This trend was more pronounced in the CVT group, supporting an association between Lcn2 induction and CVT-related astrocyte-state transitions.

At the transcriptional dynamics level, multiple inflammation- and stress-associated genes, including Ccl2, Cxcl10, Ptx3, Serpina3n, Tm4sf1, and Hmox1, were progressively upregulated along pseudotime and reached their highest expression levels at the late stages of the trajectory (Fig. 4C). The reactive gliosis marker Gfap and the metal-binding protein Mt2 exhibited similar increasing trends. These findings indicate that astrocytes progressively acquire an activated state characterized by inflammatory amplification and enhanced stress responses toward the late stages of the trajectory.

To systematically characterize this dynamic process, we further performed unsupervised clustering of pseudotime-associated genes and annotated the resulting gene modules through functional enrichment analysis (Fig. 4D, Supplementary Table 2). Multiple gene modules with distinct functional features were identified along pseudotime. A module that gradually increased toward the late stages of pseudotime (Cluster 1) was predominantly enriched in biological processes related to inflammatory responses, angiogenesis, and vascular development. By contrast, a module that was highly expressed at the early stages and subsequently declined (Cluster 2) was mainly associated with synaptic structure maintenance, cell-cell junctions, and electrophysiological regulation. In addition, an intermediate-stage module (Cluster 3) was primarily involved in protein translation and transcriptional regulation, whereas another module (Cluster 4) was enriched in receptor signaling regulation and metabolic/oxidative stress-related processes.

Collectively, the pseudotime analysis suggests that, under CVT conditions, astrocytes progress along a continuous trajectory toward a terminal Lcn2-high reactive state, accompanied by coordinated activation of inflammatory signaling and vascular-associated transcriptional programs. These findings support the interpretation that Lcn2⁺ astrocytes represent a key disease-associated state positioned at the late stage of the inferred trajectory.

### Lcn2⁺ astrocytes contribute to CVT microenvironmental remodeling through reprogrammed intercellular communication

To further investigate the system-level functional role of Lcn2⁺ astrocytes in CVT, we used CellChat to construct separate datasets containing either Lcn2⁺ or Lcn2⁻ astrocytes and comparatively analyzed their inferred cell-cell communication networks.

Global comparison revealed a greater number of inferred interactions in the dataset containing Lcn2⁺ astrocytes, whereas the overall interaction strength showed only limited changes (Fig. 5A). These findings suggest that the emergence of Lcn2⁺ astrocytes does not substantially enhance the strength of individual signaling interactions but instead reshapes intercellular information exchange by expanding the number and diversity of potential communication links.

**Figure 5.**
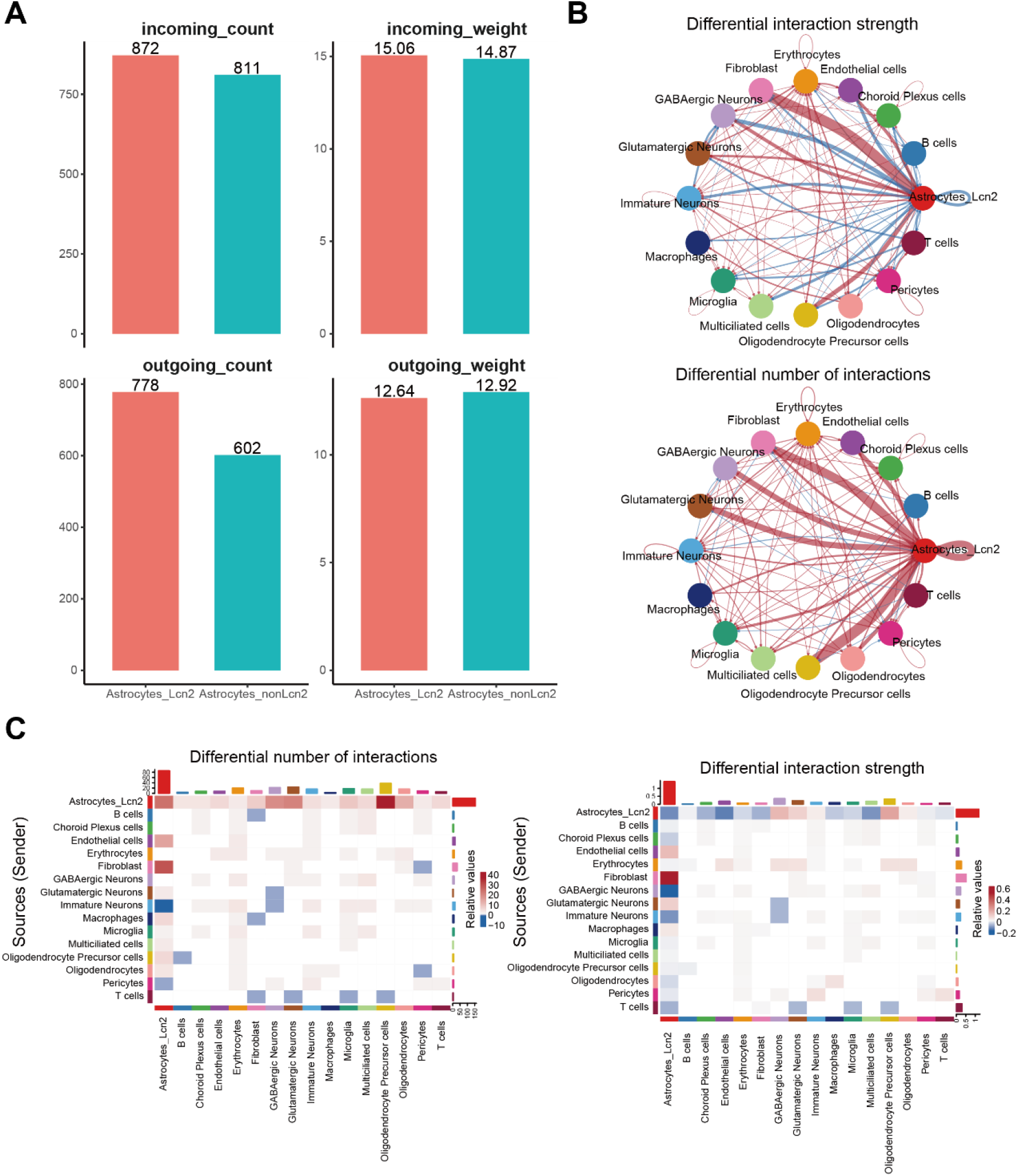
Lcn2+ astrocytes remodel the intercellular communication landscape in CVT. (A) Comparison of the total number of interactions and interaction strength between Lcn2+ astrocytes and Lcn2⁻ astrocytes. (B) Cell-cell communication networks of Lcn2+ astrocytes versus Lcn2⁻ astrocytes. (C) Differential interaction number (left) and differential interaction strength (right) associated with Lcn2+ astrocytes.

Further network visualization revealed a denser and substantially reorganized pattern of communication links in the Lcn2⁺ astrocyte-associated network compared with the network containing Lcn2⁻ astrocytes (Fig. 5B). Within this network, Lcn2⁺ astrocytes exhibited increased inferred interactions with multiple cell types, including immune cells, vascular-associated cells, and neurons, suggesting that they may function as a key communication hub within the CVT brain microenvironment.

To systematically evaluate the directionality of these alterations, we subsequently performed differential interaction analysis. In the Lcn2⁺ astrocyte-associated network, the number of inferred interactions was altered across most cell-pair combinations, whereas changes in interaction strength displayed a more selective pattern (Fig. 5C). Rather than reflecting a uniform increase or decrease in communication, these alterations represented a redistribution of signaling interactions among specific cell populations, indicating structural reprogramming of the cell-cell communication network following CVT.

Building on these findings, we further compared ligand-receptor interactions to delineate the specific signaling axes potentially mediated by Lcn2⁺ astrocytes. The predicted outgoing signals from Lcn2⁺ astrocytes were associated with several categories of biological processes. First, immune and inflammatory signaling included the MIF-(CD74+CXCR4), CSF1-CSF1R, and APP-(TREM2+TYROBP) axes. Second, vascular and barrier-regulatory signaling involved ANGPTL4- and VEGF-family pathways. In addition, neuronal and synaptic communication was represented by glutamatergic signaling through Glu-Grm5/Gria/Grik interactions and by the Tenm/Flrt-Adgrl axes (Supplementary Table 3).

Meanwhile, Lcn2⁺ astrocytes were predicted to receive diverse signaling inputs from the surrounding microenvironment, primarily originating from immune cells and vascular-associated cell populations. These incoming signals included the OSM-(LIFR/OSMR+IL6ST), SPP1-CD44/integrin, VCAM1-integrin, TNF-TNFRSF1A, and PDGF-PDGFRB axes, together with multiple extracellular matrix-related signaling pathways (Supplementary Table 3). Collectively, these predicted inputs converged on biological processes associated with inflammatory amplification and tissue injury responses.

Notably, compared with Lcn2⁻ astrocytes, Lcn2⁺ astrocytes exhibited stronger predicted communication with microglia and macrophages through ligand-receptor axes including APP-TREM2/TYROBP and APP-CD74 (Fig. 6A,B), suggesting that they may facilitate innate immune activation and inflammatory amplification. At the pathway level, Lcn2-high astrocytes showed preferential activation of multiple signaling programs compared with non-Lcn2 astrocytes, including PTN, NRXN, ADGRL, laminin, APP, VEGF, collagen, glutamate and related pathways (Fig. S3). COLLAGEN-related signaling was also markedly enhanced (Fig. 6C,D), implicating Lcn2⁺ astrocytes in extracellular matrix remodeling. In addition, increased glutamatergic signaling was observed between Lcn2⁺ astrocytes and multiple neuronal subpopulations, suggesting a potential contribution to post-stroke excitotoxicity and neural network remodeling.

**Figure 6.**
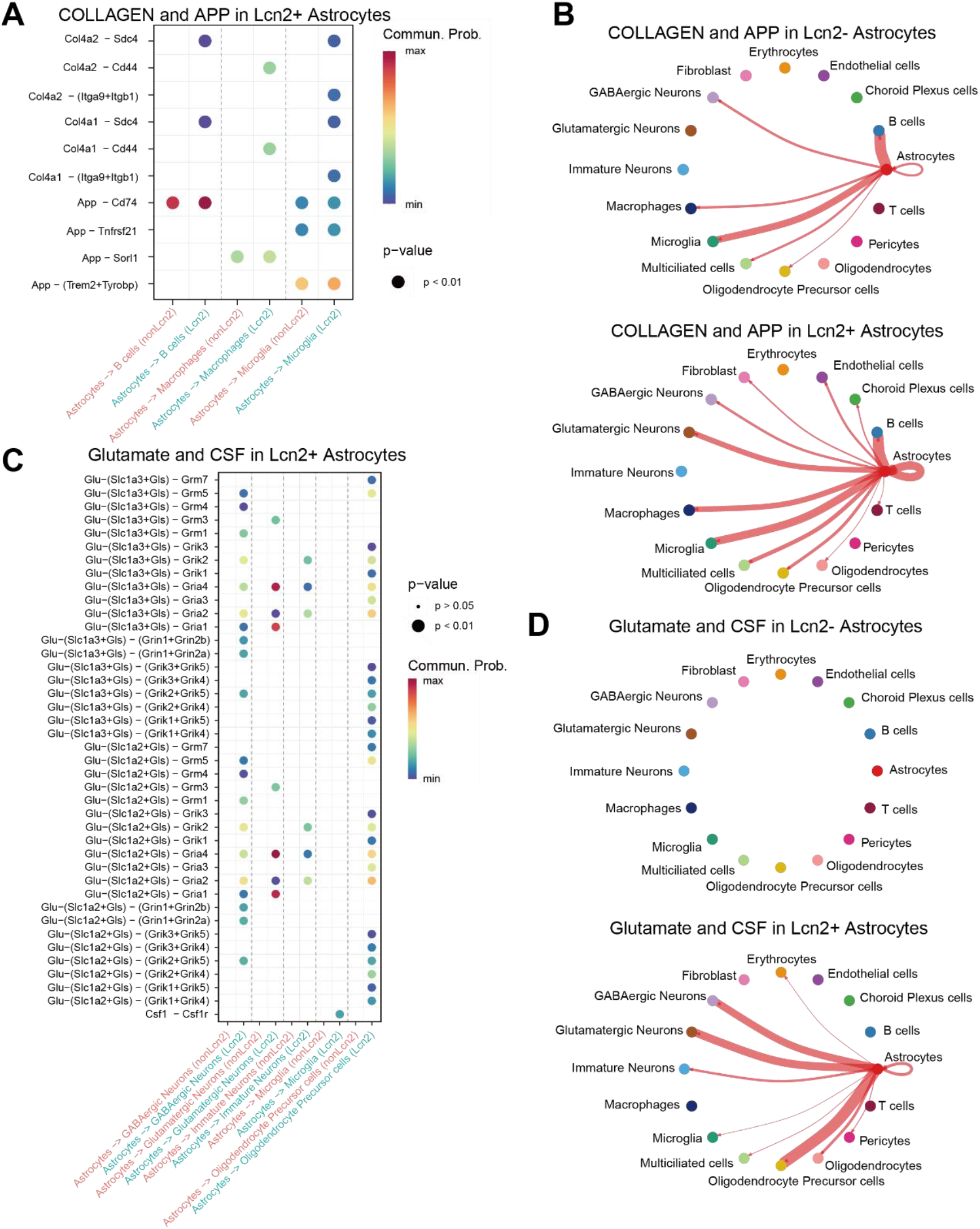
Lcn2⁺ astrocytes exhibit rewired intercellular communication networks following CVT. (A) Bubble plot showing differential ligand-receptor interactions mediated by Lcn2⁺ astrocytes compared with non-Lcn2 astrocytes. Interactions are displayed between astrocytes and recipient cell populations, including B cells, macrophages and microglia. Dot size represents statistical significance, and color intensity indicates communication probability. Notably, APP-associated signaling (APP-CD74, APP-TNFRSF21, APP-SORL1 and APP-TREM2/TYROBP) and collagen-mediated interactions were selectively enriched in Lcn2⁺ astrocytes. (B) Circular network visualization of APP and COLLAGEN signaling pathways originating from Lcn2⁺ astrocytes. Edge width corresponds to interaction strength. Lcn2⁺ astrocytes showed enhanced communication with myeloid populations, particularly macrophages and microglia, through APP- and extracellular matrix-related signaling. (C) Bubble plot displaying glutamate- and CSF-related signaling pathways associated with Lcn2⁺ astrocytes. Enhanced glutamatergic signaling was observed between Lcn2⁺ astrocytes and neuronal populations, including glutamatergic, GABAergic and immature neurons, suggesting altered neurotransmitter-associated communication following injury. (D) Circular network visualization of glutamate and CSF signaling pathways. Lcn2⁺ astrocytes exhibited strengthened interactions with neuronal and oligodendroglial lineages.

Collectively, Lcn2⁺ astrocytes appear to represent more than a population that merely responds to inflammation. Rather, they may serve as a communication hub linking the immune, vascular, and nervous systems, thereby contributing to the systematic remodeling of the CVT brain microenvironment through reprogramming of intercellular communication networks. In particular, by coordinating signaling between immune cells and neurons, Lcn2⁺ astrocytes may function as a key regulatory node connecting post-stroke neuroinflammation with neural functional remodeling and may thereby influence disease progression.

### Cross-model integrative analysis identifies Lcn2-high astrocytes following cerebrovascular occlusion, with more pronounced disease-associated features in CVT

To determine whether the Lcn2-high astrocyte state is specific to CVT, we integrated a publicly available single-cell RNA-sequencing dataset from a middle cerebral artery occlusion (MCAO) model and jointly analyzed astrocytes derived from the MCAO and CVT datasets. Following quality control, doublet removal, and cell-type annotation of the MCAO dataset, astrocytes were identified on the basis of canonical markers, including Gfap, S100b, Aldh1l1, Aqp4, Glul, and Gja1. The MCAO-derived astrocytes were then merged with the CVT astrocyte dataset, followed by Harmony-based batch correction according to data source and subsequent reclustering.

Dimensionality reduction of the integrated dataset revealed partial overlap between CVT- and MCAO-derived astrocytes, while several cellular regions remained preferentially associated with a specific model (Fig. 7A). These findings suggest that astrocytes arising under the two forms of cerebrovascular injury share a subset of transcriptional features while retaining model-dependent differences in cellular state. Further stratification by experimental group showed that control cells (S) and disease-associated cells (T and T.MCAO) occupied distinct regions within the integrated transcriptional space, with disease-associated cells markedly enriched in specific areas. This pattern indicates that injury-induced reactive astrocyte states were present in both models, represented by the CVT.3 and MCAO.0 clusters.

**Figure 7.**
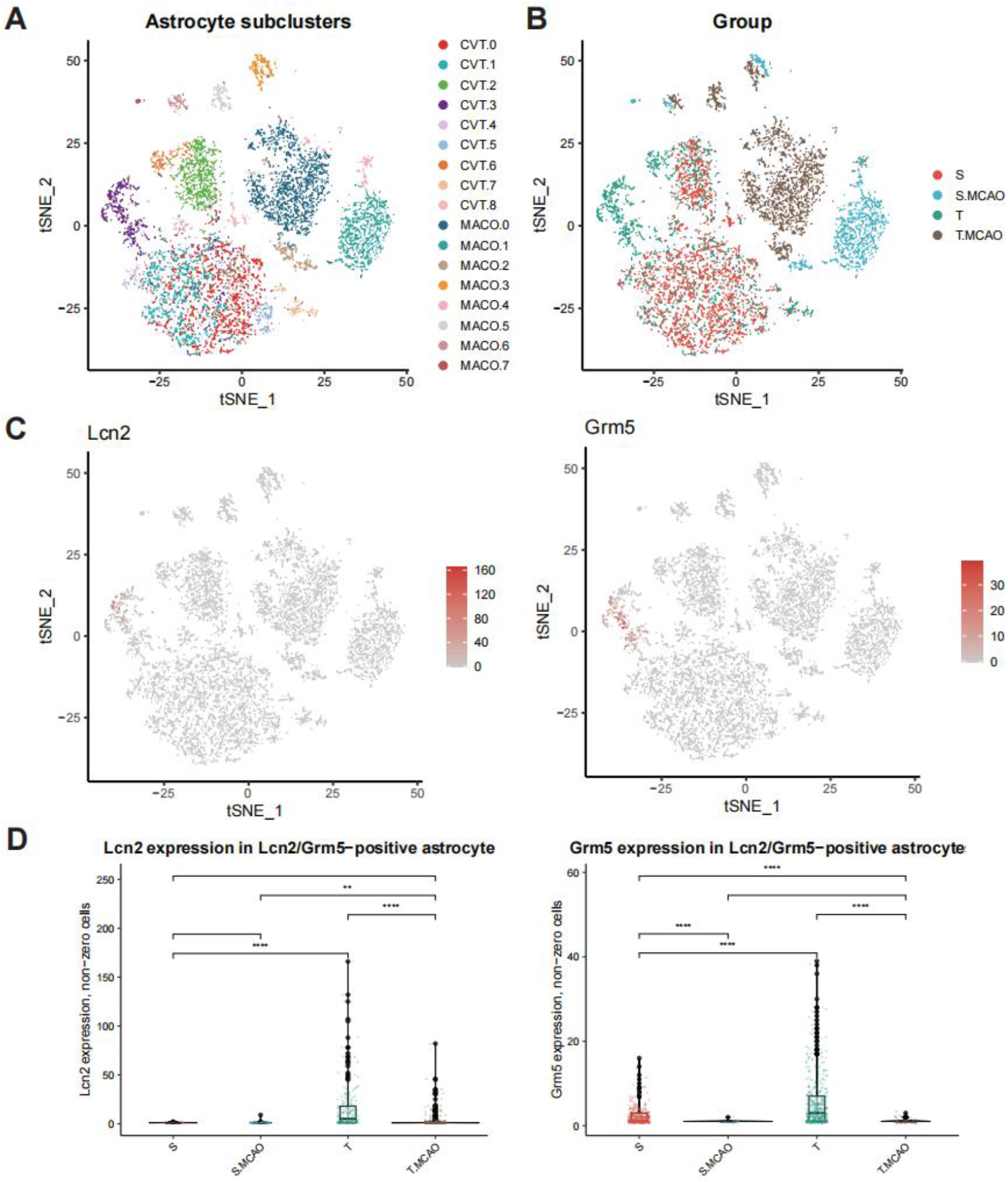
Cross-model integration of CVT and MCAO astrocytes reveals a shared but non-identical Lcn2-high reactive state. (A) t-SNE visualization of astrocytes after secondary clustering, colored by astrocyte subcluster identity. (B) t-SNE visualization of the same astrocyte compartment colored by experimental group, including S, S.MCAO, T, and T.MCAO groups. (C) Feature plots showing the distribution of Lcn2 and Grm5 expression across astrocyte subclusters. Both markers were preferentially enriched in a spatially restricted astrocyte population within the t-SNE manifold. (D) Violin plots with embedded boxplots showing Lcn2 and Grm5 expression among marker-positive astrocytes. Statistical significance was assessed using two-sided Wilcoxon rank-sum tests. p < 0.05, p < 0.01, p < 0.001, p < 0.0001.

FeaturePlot analysis further showed that Lcn2 expression was largely restricted to disease-associated cell populations and was nearly absent from the control group (Fig. 7B). Although Lcn2-high astrocytes were detectable in both models, their distributions within the integrated space did not completely overlap, suggesting that they may represent distinct reactive states under different injury contexts. Notably, Lcn2 and Grm5 expression was more prominent in CVT-associated astrocytes but comparatively weaker in the MCAO model (Fig. 7C).

Collectively, these findings indicate that Lcn2-high astrocytes are not unique to CVT but instead represent a reactive astrocyte state shared across cerebrovascular injury models. Nevertheless, their greater enrichment and stronger transcriptional activation in CVT suggest a more pronounced association with the pathological features of CVT.

### LCN2 deficiency attenuates CVT-induced brain injury in vivo

To functionally validate the role of LCN2 in secondary brain injury following CVT, we established the CVT model in wild-type (WT) and Lcn2-knockout (Lcn2-KO) mice and performed histopathological and imaging assessments of the brain at 48 h after surgery (Fig. 8A). TTC staining revealed prominent pale infarcted areas in the brains of WT-CVT mice, whereas the corresponding lesion area was markedly reduced in Lcn2-KO-CVT mice, indicating that Lcn2 deficiency attenuated the extent of CVT-induced cerebral infarction and ischemic injury (Fig. 8B). No apparent abnormal staining was observed in either the WT-sham or Lcn2-KO-sham group.

**Figure 8.**
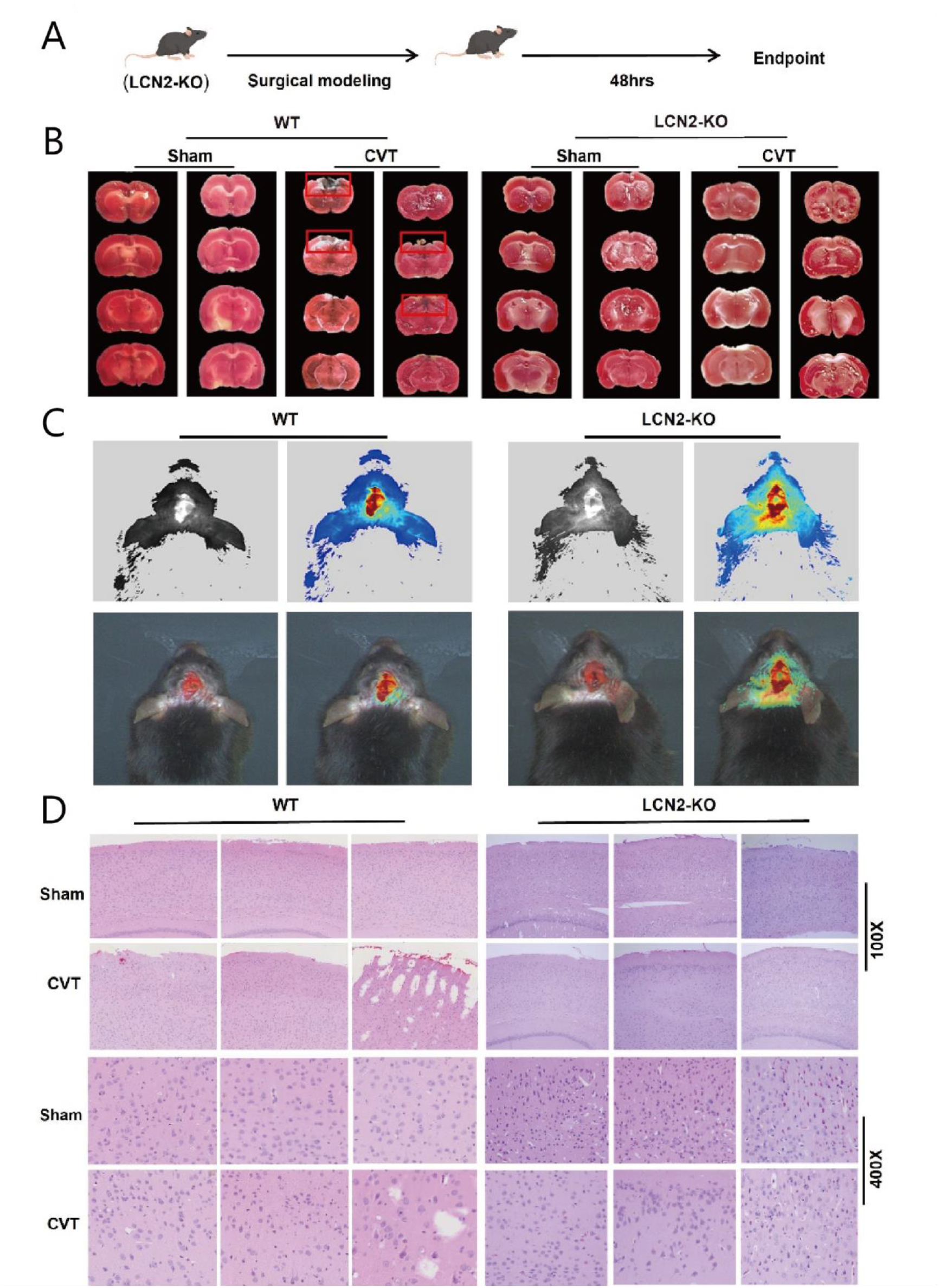
LCN2 deficiency attenuates CVT-induced brain tissue injury in vivo. (A) Schematic of the experimental design. CVT models were established in WT and LCN2-KO mice. (B) TTC staining showing a prominent infarct area in WT CVT mice, which is markedly reduced in LCN2-KO mice. (C) MRI results demonstrating that the extent of reduced cerebral blood flow and infarct regions is significantly smaller in LCN2-KO mice compared with WT mice. (D) HE staining showing attenuated brain tissue damage in LCN2-KO mice, with morphology closer to that of the sham group.

MRI analysis further supported these findings. Compared with WT-CVT mice, Lcn2-KO-CVT mice exhibited a markedly smaller area of abnormal intracerebral signal, suggesting that Lcn2 deficiency alleviated brain parenchymal injury and associated local structural and/or perfusion abnormalities following CVT (Fig. 8C). Histological analysis showed a consistent pattern. Hematoxylin and eosin staining revealed pronounced disruption of tissue architecture, disorganized cellular arrangement, and focal lesions in the WT-CVT group. By contrast, the overall brain tissue architecture was better preserved in the Lcn2-KO-CVT group, with substantially attenuated pathological damage and a morphology more closely resembling that of the sham groups (Fig. 8D). These differences were consistently observed at both low and high magnifications.

Collectively, the in vivo findings demonstrate that Lcn2 deficiency significantly attenuates CVT-induced brain tissue injury, supporting a detrimental role for LCN2 in secondary brain damage following CVT. Notably, these functional findings are consistent with the marked enrichment of Lcn2-high reactive astrocytes identified by the preceding single-cell analyses, thereby establishing a system-level association between the Lcn2-related cellular state and tissue injury.

## Methods

### Animal models and brain tissue collection

Adult mice were used to establish the cerebral venous thrombosis (CVT) model and were randomly assigned to either the CVT or sham-operated group. Cerebral venous occlusion was performed in the CVT group according to a previously described protocol with minor modifications to induce CVT. Mice in the sham group underwent the same surgical exposure and operative procedures but without venous occlusion. Brain tissues were collected at predefined postoperative time points for subsequent single-cell RNA sequencing, histological examination, and molecular analyses. For single-cell RNA sequencing, brain tissues were obtained from a total of 12 mice, including six sham-operated controls and six mice subjected to CVT.

### Brain tissue dissociation and single-cell RNA sequencing

Fresh brain tissues were rapidly dissected on ice and processed into single-cell suspensions using an enzymatic dissociation protocol optimized for neural tissues, with the aim of preserving diverse cell populations, including neurons, glial cells, vascular-associated cells, and immune cells. Briefly, the tissues were mechanically minced and enzymatically digested, followed by gentle trituration, filtration, and removal of tissue debris. Red blood cell lysis was performed when necessary. Cell viability was assessed by trypan blue exclusion, and only samples with sufficiently high viability were used for subsequent library preparation. Single-cell RNA-sequencing libraries were constructed using a droplet-based platform and sequenced on an Illumina sequencing system at sufficient depth for downstream transcriptomic analyses.

### Single-cell RNA-sequencing data preprocessing, quality control, and integration

Raw sequencing data were processed using a standard pipeline to generate gene-by-cell expression matrices. Subsequent analyses were performed in R, primarily using Seurat (version 4.3.1) for data preprocessing and integration. Low-quality cells were excluded on the basis of the number of detected genes, total unique molecular identifier (UMI) counts, and the proportion of transcripts derived from mitochondrial genes. Potential doublets were identified and removed using the CHORD tool.

Following normalization of each sample, datasets were integrated using a strategy based on shared highly variable genes to minimize technical variation while preserving biologically meaningful differences. Principal component analysis (PCA) was performed for dimensionality reduction. Cells were subsequently grouped using a graph-based unsupervised clustering algorithm constructed from the nearest-neighbor graph, and uniform manifold approximation and projection (UMAP) was used for two-dimensional visualization.

### Cell-type annotation and astrocyte subcluster analysis

Cell types in the integrated single-cell dataset were annotated according to the expression patterns of canonical marker genes. The major cell populations included astrocytes, microglia, macrophages, neurons, oligodendrocytes and oligodendrocyte precursor cells, endothelial cells, pericytes, fibroblasts, choroid plexus cells, erythrocytes, and infiltrating lymphocytes. Astrocytes were subsequently extracted from the integrated dataset and subjected to subclustering analysis to characterize their cellular heterogeneity. The resulting astrocyte subclusters were numbered according to the unsupervised clustering results, and Lcn2-enriched subclusters were identified on the basis of their Lcn2 expression patterns.

Based on the relative expression of Lcn2 across the astrocyte subclusters, clusters 9 and 10, which exhibited enriched Lcn2 expression, were operationally defined as Lcn2⁺ astrocytes, whereas the remaining astrocytes were classified as Lcn2⁻ astrocytes. For subsequent analyses, a unified grouping variable named Celltype.Lcn2.exp was generated in the metadata of the Seurat object. Cells belonging to clusters 9 and 10 were labeled as Astrocytes_Lcn2, whereas the remaining astrocytes were labeled as Astrocytes_nonLcn2.

### Functional state scoring and differential expression analysis of astrocytes

To compare the functional states of different astrocyte subclusters, single-cell pathway activity scores were calculated using Hallmark gene sets obtained from the Molecular Signatures Database (MSigDB). The analysis focused primarily on inflammation- and cytokine-related pathways, including IL-2-STAT5 signaling, IL-6-JAK-STAT3 signaling, inflammatory response, interferon-α response, and interferon-γ response. Pathway activity scores were calculated at the single-cell level and subsequently summarized according to astrocyte subcluster. Differences in pathway scores among subclusters were evaluated using appropriate nonparametric statistical tests.

Differentially expressed genes between Lcn2-high and Lcn2-low/negative astrocytes were identified to characterize differences in their transcriptional programs. Correlation analysis and heatmap visualization were further performed to illustrate the relationships among pathway activities, cellular states, and differentially expressed genes.

### Pseudotime trajectory analysis using Monocle 2

To characterize dynamic transitions in astrocyte states under CVT conditions, pseudotime trajectory analysis was performed using Monocle 2. The raw RNA count matrix was extracted from the Seurat object containing the astrocyte subset and used to construct a CellDataSet object. Gene expression was modeled using the negative binomial distribution implemented in negbinomial.size(). Size factors and gene-wise dispersions were subsequently estimated using estimateSizeFactors() and estimateDispersions(), respectively. During gene filtering, the minimum expression threshold was set to min_expr = 0.1, and only genes expressed in at least 50 cells were retained for downstream analysis.

Differential expression analyses were separately performed using differentialGeneTest() according to the experimental-group variable (mod) and pseudotime. Ordering genes used for trajectory construction were primarily selected from significantly differentially expressed genes with a q-value < 0.01. When the number of candidate genes was excessively large, the top-ranked genes were retained for trajectory inference. Dimensionality reduction was performed using the DDRTree algorithm through reduceDimension(method = “DDRTree”), after which cells were ordered along the inferred trajectory using orderCells(). To assign a biologically plausible direction to the trajectory, the root state was selected on the basis of the enrichment of control-group cells and Lcn2⁻ astrocytes within the inferred early state.

After trajectory construction, pseudotime values were extracted for individual cells. The overall trajectory and the dynamic expression patterns of selected genes along pseudotime were visualized using plot_cell_trajectory() and plot_genes_in_pseudotime(), respectively. Particular attention was given to Lcn2 and inflammation- or stress-associated genes, including Ccl2, Cxcl10, Ptx3, Serpina3n, Tm4sf1, Hmox1, Gfap, and Mt2, to assess their relationships with progressive changes in astrocyte states.

### Dynamic gene module analysis along pseudotime and functional enrichment

To characterize the transcriptional programs associated with astrocyte state transitions along pseudotime, all genes were tested for pseudotime-dependent differential expression, and genes exhibiting significant dynamic changes were identified using a q-value threshold of < 0.01. The top-ranked dynamic genes were selected to construct an expression matrix, with cells ordered according to their pseudotime values. After gene-wise Z-score normalization, unsupervised K-means clustering was applied to partition the dynamic genes into modules representing distinct temporal expression patterns.

For each gene module, Gene Ontology (GO) enrichment analysis of Biological Process (BP) terms was performed using the clusterProfiler package, with org.Mm.eg.db used as the mouse annotation database. P values were adjusted for multiple testing using the Benjamini-Hochberg method. Representative significantly enriched GO terms were selected for each module and displayed together with the expression heatmap to characterize functional programs associated with different stages of the inferred pseudotime trajectory. Heatmaps were generated from the standardized expression matrix after ordering cells by pseudotime. The ComplexHeatmap package was used to visualize dynamic gene modules, pseudotime gradient annotations, and the corresponding functional annotations.

### Cell-cell communication analysis using CellChat

To evaluate differences in intercellular communication between Lcn2-high and Lcn2-low/negative astrocytes within the brain microenvironment, ligand-receptor interactions were inferred using CellChat. Briefly, CellChat objects were generated using createCellChat(), and the mouse ligand-receptor database CellChatDB.mouse was assigned for subsequent analysis. Overexpressed genes and ligand-receptor interactions were identified, after which cell-cell communication probabilities and pathway-level communication probabilities were computed. The inferred communication networks were then aggregated, and network centrality measures were calculated to characterize the relative signaling roles of individual cell populations.

For comparative analyses, the overall number and strength of inferred interactions involving Lcn2-high or Lcn2-low/negative astrocytes were evaluated using compareInteractions(). Differential networks representing changes in interaction number and interaction strength were visualized using netVisual_diffInteraction(). Major ligand-receptor axes were further identified by separately considering Lcn2-high and Lcn2-low/negative astrocytes as signal senders or receivers. Heatmaps were used to display differences in the inferred number and strength of interactions between astrocytes and other cell populations.

### Cross-model integration analysis of astrocytes from CVT and MCAO

To compare the similarities and differences in astrocyte states between CVT and arterial cerebral ischemia, a publicly available single-cell RNA-sequencing dataset derived from a middle cerebral artery occlusion (MCAO) model was included.

Following data preprocessing, quality control, and doublet removal, astrocytes were identified and extracted on the basis of canonical marker genes, including Gfap, S100b, Aldh1l1, Aqp4, Glul, and Gja1. MCAO-derived astrocytes were subsequently merged with CVT-derived astrocytes. The combined dataset was normalized using SCTransform, and batch effects associated with data source (CVT or MCAO) were corrected using Harmony. Dimensionality reduction, unsupervised clustering, and visualization were then repeated on the integrated astrocyte dataset.

Within the integrated object, the expression distributions of Lcn2 and Grm5 across the two models were visualized using FeaturePlot() and VlnPlot(). In addition, the proportion of Lcn2-high astrocytes and the mean Lcn2 expression level were calculated for each experimental group (S, T, S.MCAO, and T.MCAO) to assess the shared and model-dependent characteristics of the Lcn2-high astrocyte state in CVT and MCAO.

### In vivo functional validation of LCN2 in WT and Lcn2-KO mice subjected to CVT

To investigate the functional role of LCN2 in secondary brain injury following CVT, the CVT model was established in wild-type (WT) and Lcn2-knockout (Lcn2-KO) mice, with the corresponding sham-operated animals included as controls. The animals were assigned to four groups: WT-sham, WT-CVT, Lcn2-KO-sham, and Lcn2-KO-CVT. Successful deletion of Lcn2 in the knockout mice was confirmed by genotyping before the animals were used for CVT modeling and subsequent in vivo functional assessments.

At predefined postoperative time points, mice were euthanized, and the brains were rapidly collected and sectioned coronally. Brain sections were subjected to 2,3,5-triphenyltetrazolium chloride (TTC) staining to evaluate the extent of cerebral ischemic or infarct-related injury. MRI was performed at the designated postoperative time point to further assess brain lesions and abnormal intracerebral signals in WT and Lcn2-KO mice following CVT. MRI findings were compared between the two genotypes to determine the effect of Lcn2 deficiency on CVT-induced brain tissue injury.

For histopathological examination, brain tissues were fixed, embedded, sectioned, and stained with hematoxylin and eosin. Tissue architecture, cellular disorganization, focal lesions, and other pathological alterations were evaluated under both low and high magnifications.

### Statistical analysis and data visualization

All statistical analyses were performed in R. Comparisons of cell-type proportions, pathway activity scores, module scores, gene expression levels, and quantitative behavioral or histological measurements between two independent groups were conducted using the two-sided Wilcoxon rank-sum test, unless otherwise specified. A P value of < 0.05 was considered statistically significant. Unless otherwise indicated, data visualization was performed primarily using ggplot2, ComplexHeatmap, Monocle 2, CellChat, and functions implemented in Seurat.

## Discussion

Cerebral venous thrombosis (CVT) is a cerebrovascular disorder characterized by distinct hemodynamic features, and its pathological processes differ from those of conventional arterial ischemic stroke. Impaired venous outflow and elevated venous pressure can lead to cerebral venous congestion, blood-brain barrier disruption, cerebral edema, and hemorrhagic infarction [25]. However, how different cell types respond to the abnormal vascular environment during secondary brain injury following CVT, and which cellular states participate in disease progression, remain poorly understood at single-cell resolution.

In the present study, single-cell RNA sequencing was used to systematically characterize changes in the cellular composition and functional states of brain tissue following CVT. We found that CVT did not induce a large-scale expansion of the astrocyte population but instead predominantly remodeled astrocyte functional states. Within the astrocyte compartment, we identified a disease-associated reactive state characterized by high Lcn2 expression, hereafter referred to as the Lcn2-high astrocyte state. This state was markedly enriched under CVT conditions and exhibited enhanced inflammatory responses, cytokine signaling, and activation of stress-related programs associated with iron homeostasis. Further cross-model analyses and validation in Lcn2-deficient mice revealed a consistent biological association between the Lcn2-related cellular state and CVT-induced brain tissue injury. Collectively, these findings support a new perspective in which the astrocyte response following CVT is not simply characterized by an increase in cell abundance but instead involves the remodeling of specific functional states that contribute to the formation of the disease-associated microenvironment.

Previous studies of glial responses and cellular heterogeneity following cerebrovascular injury have focused primarily on models of arterial ischemia, such as middle cerebral artery occlusion (MCAO) [26]. However, CVT has a pathological basis distinct from that of arterial occlusion, including persistent venous hypertension, tissue congestion, increased vascular permeability, and a more pronounced tendency toward hemorrhage [25]. Therefore, the local microenvironment generated by CVT may induce cellular response patterns that differ from those observed in conventional ischemic stroke.

One of the important findings of the present study was that, although CVT resulted in inflammatory cell infiltration and marked alterations in the brain microenvironment, the overall proportion of astrocytes did not increase significantly. This finding suggests that the astrocyte response following CVT does not primarily involve the expansion of reactive glial cells in the conventional sense, but instead reflects a reorganization of cellular states and functional programs.

In recent years, single-cell and spatial omics studies have progressively changed the traditional understanding of reactive astrocytes. Growing evidence indicates that astrocytes are not a single homogeneous population but can adopt diverse functional states depending on the type of injury, disease stage, and local microenvironment [17,26,27]. Some of these states may contribute to the restriction of inflammation, stabilization of neural networks, and tissue repair, whereas others may be associated with the amplification of inflammation, disruption of the blood-brain barrier, and neurovascular dysfunction [17,27–30]. The Lcn2-high astrocyte state identified in the present study further supports this concept, indicating that CVT does not induce all astrocytes to enter a uniform activation state but instead selectively enriches a reactive subpopulation with distinct molecular characteristics.

LCN2 is an important stress-inducible secreted protein that has been implicated in the regulation of inflammation, iron homeostasis, and oxidative stress responses in various forms of central nervous system injury [22,31]. However, previous studies have mainly focused on the function of LCN2 as an individual molecular factor, whereas whether its expression represents a distinct cellular state has not been sufficiently characterized.

Our single-cell analysis revealed that Lcn2 expression was not uniformly distributed across all astrocytes but was predominantly concentrated within specific astrocyte subpopulations. This Lcn2-high state was accompanied by enhanced IL-6-JAK-STAT3 signaling, interferon-response programs, inflammatory responses, and NF-κB-related transcriptional programs, suggesting that it represents a highly stress-associated reactive astrocyte state.

Importantly, the identification of Lcn2-high astrocytes does not imply that all reactive astrocytes exert detrimental effects. Recent studies have demonstrated that reactive astrocytes are highly context dependent, with their functions being determined by disease type, stage of injury, and the local cellular microenvironment [17,27–29]. Therefore, rather than simply classifying Lcn2-high astrocytes as “harmful astrocytes,” we interpret them as an inflammatory reactive state associated with CVT.

In addition to inflammation-related signaling, we found that Lcn2-high astrocytes were enriched for ferroptosis-related gene programs and expressed molecules associated with metal binding and oxidative stress, including Hmox1 and Mt2. Previous studies have shown that LCN2 participates in the regulation of iron homeostasis and redox balance and, in the context of stroke, may promote lipid peroxidation, ferroptosis, and blood-brain barrier injury in endothelial cells [22,31].

However, it is important to emphasize that the transcriptomic enrichment observed in the present study does not demonstrate that Lcn2-high astrocytes themselves undergo ferroptosis. A more reasonable interpretation is that Lcn2-high astrocytes exist in a cellular state characterized by disrupted iron homeostasis and increased oxidative stress. Through alterations in the local inflammatory environment, paracrine signaling, and homeostasis of the neurovascular unit, this state may influence the susceptibility of neighboring cells to injury-related stimuli.

Given that CVT is characterized by pronounced venous congestion, blood-brain barrier disruption, and a tendency toward hemorrhagic transformation [25], this Lcn2-associated inflammatory-oxidative stress state may represent one of the important pathological features distinguishing CVT from conventional arterial ischemia.

Cell-cell communication analysis further suggested that Lcn2-high astrocytes participate in a more complex network of potential interactions with immune cells, vascular-associated cells, and neurons. Previous studies have shown that reactive astrocytes may contribute to vascular repair, extracellular matrix remodeling, stabilization of neural networks, and regulation of the blood-brain barrier following stroke [26,28–30]. In particular, APP-related signaling, extracellular matrix-associated signaling, and neurotransmitter-related pathways were more prominent in the Lcn2-high state. These findings suggest that this astrocyte state may participate in coordinating inflammatory responses, alterations in the vascular microenvironment, and the regulation of neuronal homeostasis.

However, it should be emphasized that cell-cell communication analyses inferred from ligand-receptor expression profiles inherently reflect potential signaling relationships rather than experimentally confirmed physical interactions or functional signal transmission. Moreover, different computational algorithms and ligand-receptor databases may generate substantially different predictions [32,33]. Therefore, the cell-cell communication results presented in this study should primarily be interpreted as candidate regulatory networks that provide directions for subsequent validation using spatial omics and functional experiments.

Although different forms of cerebrovascular injury have distinct pathological foundations, they can all induce hypoxia, inflammation, and glial responses [17,26,27]. An important question is therefore whether the Lcn2-high astrocyte state is specific to CVT. By comparing our data with a publicly available single-cell RNA-sequencing dataset from an MCAO model, we found that the Lcn2-high state was not entirely specific to CVT but was detectable across different models of cerebrovascular injury. This finding suggests that Lcn2-high astrocytes may represent a shared cellular response program following cerebrovascular injury.

Nevertheless, marked differences remained between the two models. The venous congestion, sustained elevation of vascular pressure, and blood-brain barrier disruption characteristic of CVT [25] may further promote the emergence or enhancement of this reactive state. Therefore, rather than considering Lcn2-high astrocytes to be a CVT-specific subpopulation, they may be more appropriately interpreted as an injury-associated cellular state that is shaped by the disease microenvironment and is more prominently manifested in CVT.

To further evaluate the role of LCN2 in CVT-induced injury, we performed in vivo validation using Lcn2-deficient mice. The results showed that Lcn2 deficiency markedly attenuated CVT-induced tissue injury, as evidenced by a reduced infarct area, a smaller region of abnormal imaging signals, and improved tissue architecture. These findings provide whole-animal evidence supporting the involvement of LCN2 in secondary brain injury following CVT.

Previous studies of stroke have also suggested that LCN2 expression in reactive astrocytes is associated with neuroinflammation and neurodegenerative injury, and that LCN2 can additionally act on cerebrovascular endothelial cells and influence blood-brain barrier homeostasis [22,31]. However, because the present study used a whole-body Lcn2-knockout model, these results cannot distinguish the relative contributions of LCN2 derived from different cell types. In addition to astrocytes, vascular-associated cells may also participate in LCN2-related injury responses [22,31].

Therefore, the functional validation performed in this study supports LCN2 as a pathological factor associated with CVT, but future studies using astrocyte-specific genetic models will be required to determine the direct causal contribution of Lcn2-high astrocytes.

Several limitations of this study should be acknowledged. First, the present investigation was based on a mouse model of CVT, and the extent to which the Lcn2-high astrocyte state is conserved in human CVT remains to be determined. Second, single-cell trajectory and cell-cell communication analyses primarily provide information regarding the continuity of cellular states and potential intercellular interactions, respectively, and cannot substitute for real-time lineage tracing or functional intervention experiments. In particular, ligand-receptor-based communication inference may be influenced by the computational algorithm, the ligand-receptor database used, and the detection efficiency of gene expression [32,33]. Third, although the Lcn2-deficient model demonstrated the overall importance of LCN2, it could not fully resolve the specific contribution of astrocyte-derived LCN2. Future studies integrating spatial transcriptomics, human CVT samples, and cell type-specific genetic manipulation will be required to further define the precise functional role of Lcn2-high astrocytes in the pathological progression of CVT.

In summary, single-cell transcriptomic analysis revealed extensive remodeling of astrocyte states in the brain following CVT and identified a disease-associated reactive state characterized by high Lcn2 expression. This state exhibited enhanced inflammatory responses, stress programs related to iron homeostasis, and features associated with remodeling of the neurovascular-immune microenvironment, while also showing partial conservation across different models of cerebrovascular injury. In parallel, Lcn2 deficiency attenuated CVT-associated tissue injury, supporting a pathological role for LCN2 at the whole-organism level.

These findings suggest that brain injury following CVT arises not only from vascular occlusion itself but is also accompanied by the dynamic reorganization of specific glial cell states. The Lcn2-high astrocyte state may represent an important cellular program linking inflammation, oxidative stress, and alterations in the neurovascular microenvironment, thereby providing a new cellular framework for understanding secondary brain injury following CVT.

## Declarations

## Ethical Approval and Consent to participate

All animal experimental procedures were reviewed and approved by the Institutional Animal Care and Use Committee of Kangtai Medical Laboratory Services (Hebei) Co., Ltd. (approval No. MDL2024-12-28-01; approved on 28 December 2024). All procedures were conducted in accordance with the approved protocol and relevant institutional guidelines for the care and use of laboratory animals. Consent to participate was not applicable because no human participants were involved in this study.

## Consent for publication

Not applicable.

## Availability of data and materials

The single-cell RNA-sequencing data generated in this study have been deposited in the NCBI Gene Expression Omnibus (GEO) under submission number SUB16606718. The corresponding GSE accession number will be released upon publication. All other datasets analysed during the current study are available from the corresponding author on reasonable request.

## Competing interests

The authors declare that they have no competing interests.

## Funding

This work was supported by the General Program of the Medical Science and Technology Research Fund of Guangdong Province (No. A2024393), the General Program of the Shenzhen Natural Science Foundation (No. JCYJ20230807115312026), the Major Science and Technology Project of the 2023 Nanshan District Health System (No. NSZD2023046), the Guangdong-Shenzhen Joint Fund of the Guangdong Basic and Applied Basic Research Foundation (No. 2023A1515110996), and the Shenzhen Science and Technology Innovation Program of the Shenzhen Municipal Science and Technology Innovation Council (No. JCYJ20250604183519025).

## Authors’ contributions

L.K. (Ling Kui) contributed to conceptualization, study design, investigation planning, and writing of the original manuscript. S.-W.Z. (Si-Wei Zhou) conducted the literature investigation and preliminary research and participated in manuscript review and editing. Y.-M.J. (Yin-Ming Jiao) performed the animal experiments and experimental procedures and acquired the data. R.Z. (Rong Zhang) contributed to manuscript review, refinement, and language editing. Y.-Q.Z. (Yan-Qing Zhou) and L.-B.L.(Ling-Bin Liu) coordinated the research, managed resources, and provided computational resources. G.-Y.W. (Guo-Yun Wang) contributed to manuscript review and critical revision and assisted with manuscript preparation. K.-X.X. (Ke-Xu Xiong) performed data and bioinformatics analyses, conducted data visualization, prepared the figures, and revised the manuscript. All authors reviewed and approved the final manuscript.

## Acknowledgements

Not applicable.

## Notes

### Competing Interest Statement

The authors have declared no competing interest.

