## Supplementary Fig. 1. for "LCN2+ Astrocytes Mark a Pro-inflammatory Astrocyte State Associated with Secondary Brain Injury After Cerebral Venous Thrombosis"

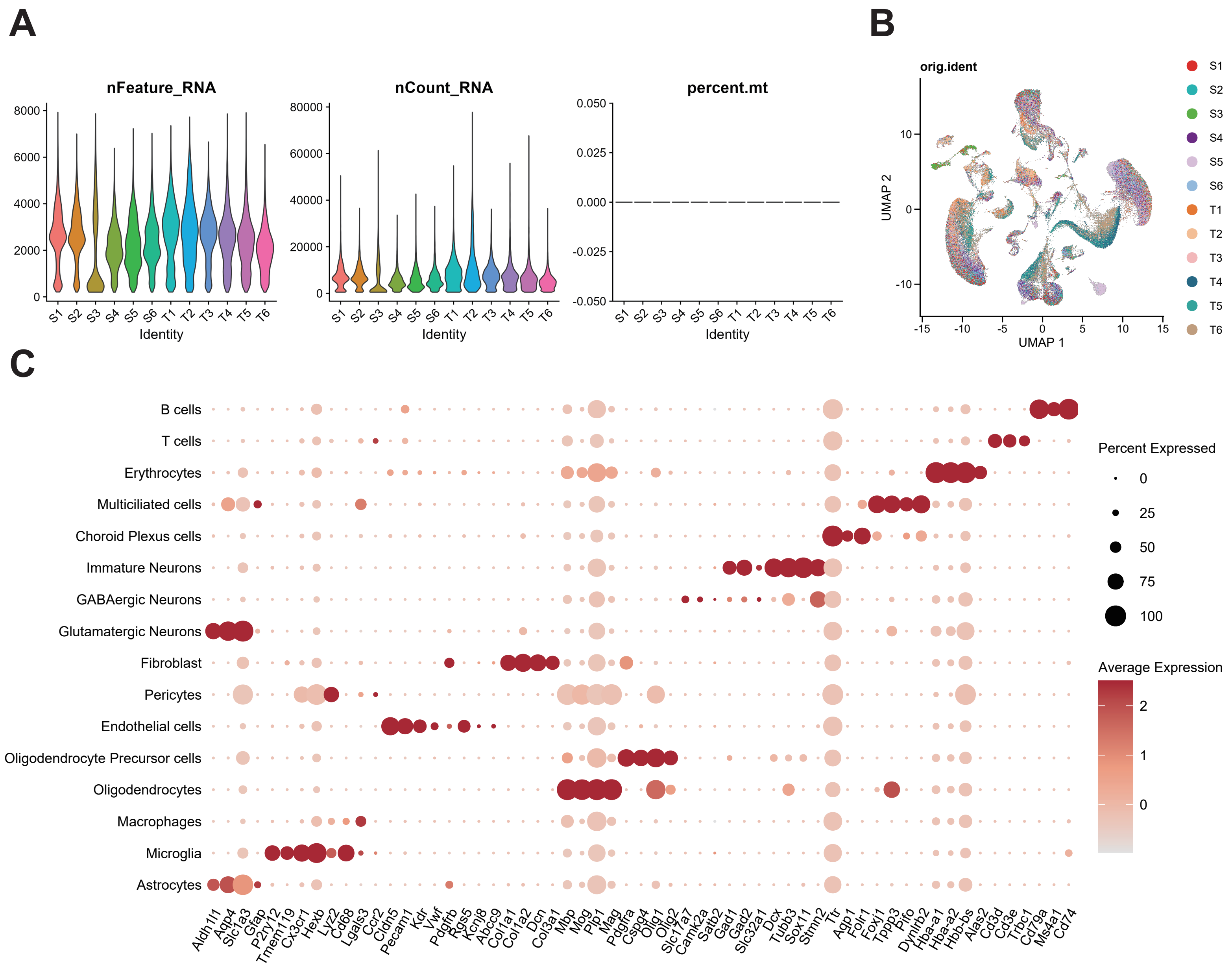


**Supplementary Figure 1. Single-cell RNA sequencing analysis identifies major cell populations in the CVT mouse brain.**

**(A)** Quality control assessment of single-cell RNA sequencing data.

**(B)** Uniform manifold approximation and projection (UMAP) visualization of integrated single-cell transcriptomic profiles from sham (S1–S6) and CVT (T1–T6) mouse brains.

**(C)** Dot plot showing canonical marker gene expression used for annotation of major brain cell populations.
