## Supplementary Fig. 2. for "LCN2+ Astrocytes Mark a Pro-inflammatory Astrocyte State Associated with Secondary Brain Injury After Cerebral Venous Thrombosis"

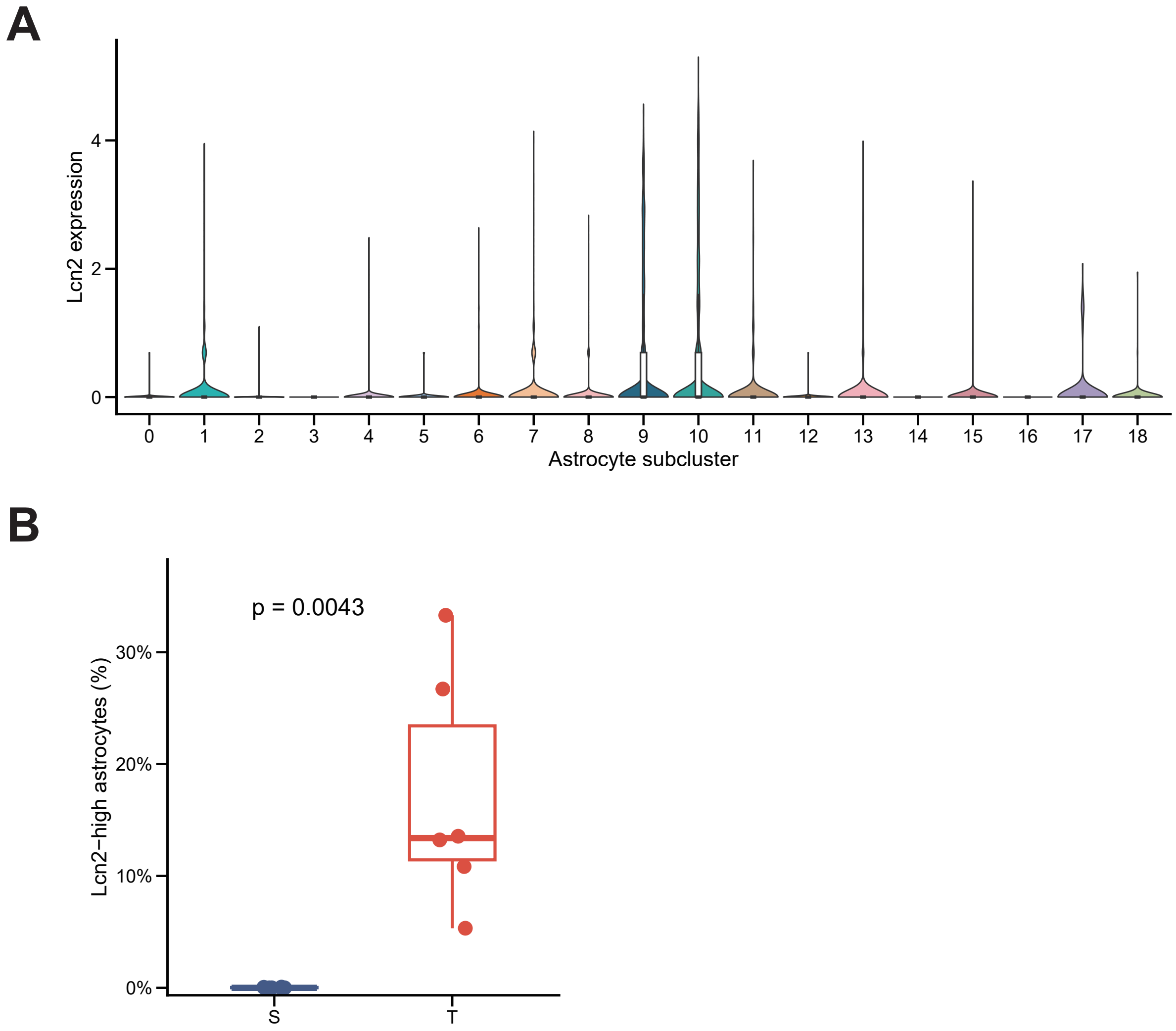


**Supplementary Figure 2. Identification of a CVT-associated Lcn2-high astrocyte population.**

**(A)** Violin plots showing Lcn2 expression across astrocyte subclusters identified from single-cell transcriptomic analysis. Lcn2 expression was markedly enriched in a specific astrocyte subpopulation, corresponding to the Lcn2-high astrocyte state.

**(B)** Quantification of the proportion of Lcn2-high astrocytes among total astrocytes in sham and CVT conditions. CVT induction significantly increased the abundance of Lcn2-high astrocytes compared with sham controls (P = 0.0043).
